# Canu: scalable and accurate long-read assembly via adaptive *k*-mer weighting and repeat separation

**DOI:** 10.1101/071282

**Authors:** Sergey Koren, Brian P. Walenz, Konstantin Berlin, Jason R. Miller, Nicholas H. Bergman, Adam M. Phillippy

**Affiliations:** Genome Informatics Section, Computational and Statistical Genomics Branch, National Human Genome Research Institute, National Institutes of Health, Bethesda, MD USA; Invincea Labs, Arlington, VA USA; J. Craig Venter Institute, Rockville, MD USA; National Biodefense Analysis and Countermeasures Center, Frederick, Maryland USA

**Author notes:** These authors contributed equally to this work (Sergey Koren, Brian P. Walenz).

**Keywords:** de novo assembly, single-molecule sequencing, nanopore sequencing

## Abstract

Long-read single-molecule sequencing has revolutionized *de novo* genome assembly and enabled the automated reconstruction of reference-quality genomes. However, given the relatively high error rates of such technologies, efficient and accurate assembly of large repeats and closely related haplotypes remains challenging. We address these issues with Canu, a successor of Celera Assembler that is specifically designed for noisy single-molecule sequences. Canu introduces support for nanopore sequencing, halves depth-of-coverage requirements, and improves assembly continuity while simultaneously reducing runtime by an order of magnitude on large genomes versus Celera Assembler 8.2. These advances result from new overlapping and assembly algorithms, including an adaptive overlapping strategy based on *tf-idf* weighted MinHash and a sparse assembly graph construction that avoids collapsing diverged repeats and haplotypes. We demonstrate that Canu can reliably assemble complete microbial genomes and near-complete eukaryotic chromosomes using either PacBio or Oxford Nanopore technologies, and achieves a contig NG50 of greater than 21 Mbp on both human and *Drosophila melanogaster* PacBio datasets. For assembly structures that cannot be linearly represented, Canu provides graph-based assembly outputs in graphical fragment assembly (GFA) format for analysis or integration with complementary phasing and scaffolding techniques. The combination of such highly resolved assembly graphs with long-range scaffolding information promises the complete and automated assembly of complex genomes.

## Introduction

The goal of genome assembly is to reconstruct a complete genome from many comparatively short sequencing reads. Overlapping reads that originate from the same region of the genome can be joined together to form contigs, but genomic repeats longer than the overlap length lead to ambiguous reconstructions and fragment the assembly (Phillippy et al. 2008; Nagarajan and Pop 2009). There are two strategies for overcoming this fundamental limitation—increasing the effective read length, and separating non-exact repeats based on copy-specific variations. Recently, single-molecule sequencing has revolutionized assembly by producing reads longer than 10 kbp (Gordon et al. 2016), which has significantly reduced the number of unresolvable repeats (Koren et al. 2012) and enabled the complete assembly of microbial genomes (Chin et al. 2013; Koren et al. 2013; Koren and Phillippy 2014). These long reads also aid assembly phasing (Chin et al. 2016), where the conserved alleles in a diploid, polyploid, or meta-genome can be thought of as a special kind of repeat. However, in contrast to improved read length, single-molecule sequencing is less accurate than past technologies (Eid et al. 2009; Schneider and Dekker 2012), requiring sensitive alignment methods and limiting the discrimination of divergent alleles and non-exact repeats. Nevertheless, PacBio single-molecule real-time (SMRT) sequencing exhibits a largely unbiased and random error model (Ross et al. 2013), enabling assemblies that exceed short-read data both in terms of quality and continuity (Chin et al. 2013; Koren et al. 2013). Oxford Nanopore strand sequencing can also produce highly continuous assemblies, but current biases in base calling prohibit an accurate consensus sequence without the addition of complementary data (Loman et al. 2015).

The increased read length and error rate of single-molecule sequencing has challenged genome assembly programs originally designed for shorter, highly accurate reads. Several new approaches have been developed to address this, roughly categorized as hybrid, hierarchical, or direct. See (Koren and Phillippy 2014) for a review. Hybrid methods use single-molecule reads to reconstruct the long-range structure of the genome, but rely on complementary short reads for accurate base calls (Koren et al. 2012; Hackl et al. 2014; Lee et al. 2014; Salmela and Rivals 2014; Antipov et al. 2016; Ye et al. 2016). Hierarchical methods do not require a secondary technology and instead use multiple rounds of read overlapping (alignment) and correction to improve the quality of the single-molecule reads prior to assembly (Chin et al. 2013; Koren et al. 2013). Finally, direct methods attempt to assemble single-molecule reads from a single overlapping step without any prior correction (Li 2016; Tørresen et al. 2016). All three approaches are capable of producing an accurate final assembly. However, our goal is the complete reconstruction of entire genomes, so we focus here on the hierarchical strategy because it has produced the most continuous *de novo* assemblies to date (Berlin et al. 2015; Chakraborty et al. 2016).

## Results

Canu is a new single-molecule sequence assembler that improves upon and supersedes the now unsupported Celera Assembler (Myers et al. 2000; Miller et al. 2008). Recently, we introduced the MinHash Alignment Process (MHAP) to overcome the computational bottleneck of overlapping noisy, single-molecule sequencing reads (Berlin et al. 2015). Combining this technique with PBcR (Koren et al. 2012) and Celera Assembler, we demonstrated near-complete eukaryotic assemblies from PacBio sequencing alone (Berlin et al. 2015). Building on this work, we developed Canu to (1) integrate our methods into a single, comprehensive assembler, (2) support both PacBio and Oxford Nanopore data, (3) lower runtime and coverage requirements, and (4) improve repeat and haplotype separation. As a result, Canu improves runtime by an order of magnitude for mammalian genomes and outperforms hybrid methods with as little as 20× single-molecule coverage. At higher coverage, reference-quality *de novo* assemblies are possible, including the complete assembly of euchromatic chromosomes from either PacBio or Nanopore sequencing. In addition, Canu's improved graph construction algorithm separates closely related repeats and alleles based on a statistical model of read error, which will be important for future work on diploid, polypoloid, and metagenomic assembly. Canu source code and pre-compiled binaries are freely available under a GPLv2 license from https://github.com/marbl/canu.

### Architecture

To improve usability and performance on single-molecule sequence data, Canu introduces several novel features including computational resource discovery, adaptive *k*-mer weighting, automated error rate estimation, sparse graph construction, and graphical fragment assembly (GFA) (Li 2016) outputs. The Canu pipeline consists of three stages—correction, trimming, and assembly (Figure 1)—each of which can run independently or in series (e.g. only read correction, or assembly without correction, etc.). When running in a parallel environment, Canu will auto-detect available resources and configure itself to maximize resource utilization. It is currently the most efficient single-molecule read assembler available for large genomes, requiring approximately 20,000 CPU hours to assemble a human genome, compared to ~60,000 required for Falcon (Chin et al. 2016) and >250,000 required for Celera Assembler v8.2 (Berlin et al. 2015). In addition to these runtime improvements, the resulting assemblies are significantly more continuous than prior versions.

**Figure 1.**
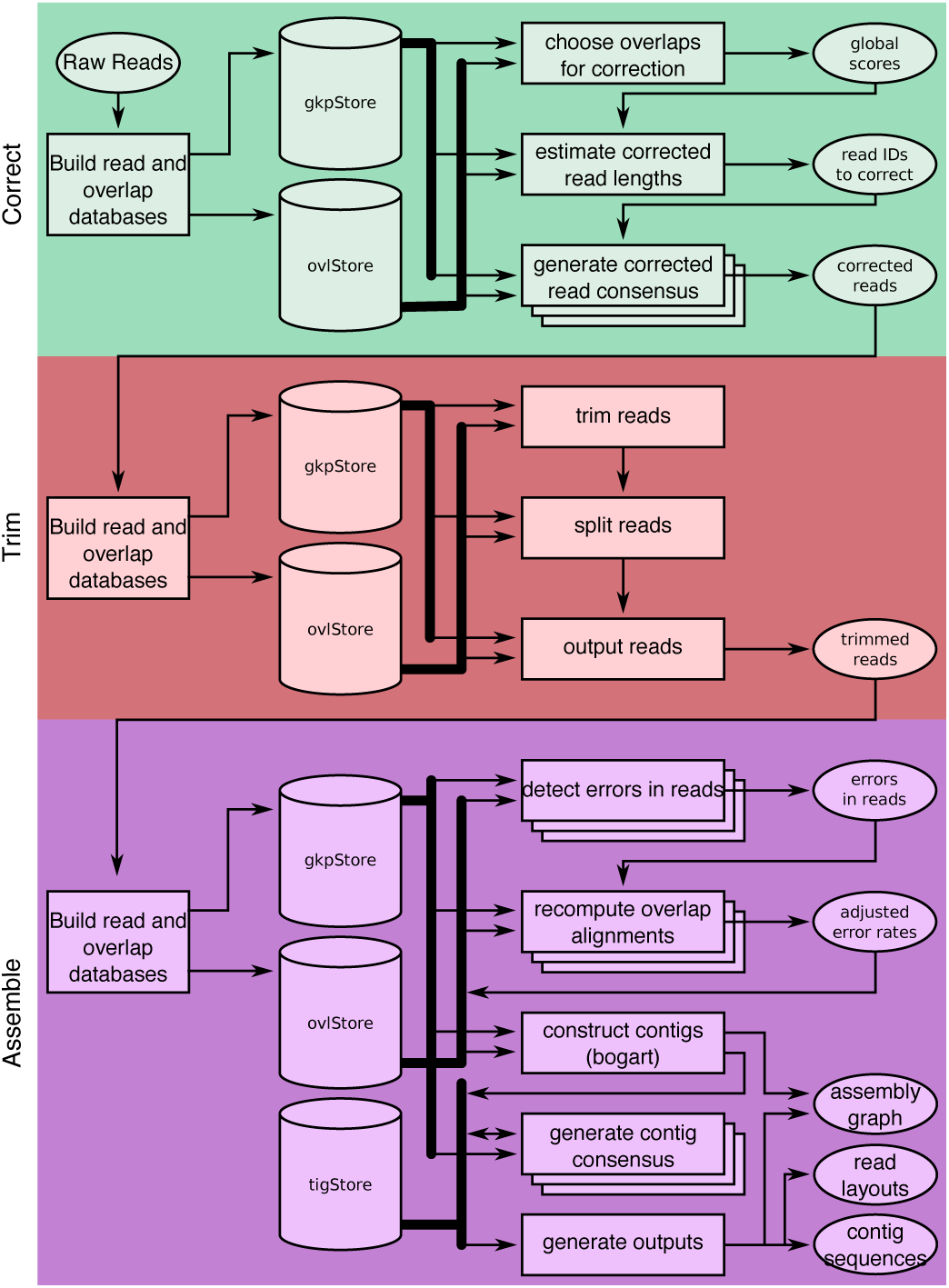
**A full Canu run includes three stages: correction (green), trimming (red), and assembly (purple)**. Canu stages share an interface for binary on-disk stores (databases) as well as parallel store construction. In all stages, the first step constructs an indexed store of input sequences, generates a *k*-mer histogram, constructs an indexed store of all-vs-all overlaps, and collates summary statistics. The correction stage (green) selects the best overlaps to use for correction, estimates corrected read lengths, and generates corrected reads. The trimming stage (red) identifies unsupported regions in the input and trims or splits reads to their longest supported range. The assembly stage (purple) makes a final pass to identify sequencing errors; constructs the best overlap graph; and outputs contigs, an assembly graph, and summary statistics.

### Adaptive MinHash *k*-mer weighting

Optimal handling of repeats is a challenge, because in addition to fragmenting assemblies, repeats also cause computational bottlenecks during overlapping. Read overlapping typically proceeds in two stages, first building a list of read pairs that share some similarity and then performing a more direct comparison of those read pairs (e.g. dynamic programming) (Sutton et al. 1995). Candidate overlaps are typically found in the first stage by identifying shared *k*-mers (length *k* substrings) between all pairs of reads. However, repeats reduce the entropy of the *k*-mer distribution compared to random sequence, and the frequent occurrence of some *k*-mers significantly increases the number of candidate overlaps that must be processed by the more expensive second stage. A common solution is to mask low-complexity sequence or ignore highly repetitive *k*-mers during indexing (Ning et al. 2001), as is done by many assemblers including Celera Assembler (Myers et al. 2000), Falcon (Chin et al. 2016), and Miniasm (Li 2016). However, depending on how many repeating *k*-mers are ignored, some fraction of correct overlaps will not be detected.

Canu takes a more resilient approach to handling repeats that probabilistically reduces, but does not eliminate, the chance a repetitive *k*-mer will be selected for overlapping. This weighting is achieved via a MinHash overlapping strategy. Rather than comparing individual *k*-mers to identify potential read overlaps, Canu uses the previously described MinHash Alignment Process (MHAP) to compare compressed sketches of entire reads (Berlin et al. 2015). Because each MinHash sketch contains a fixed-size subset of *k*-mers selected from a read, the probability of including particular *k*-mers in a sketch can be adjusted. For instance, a repetitive *k*-mer occurring many times throughout the genome should have a reduced weight, because it carries relatively little information regarding the origin of the read. In contrast, a relatively unique *k*-mer occurring multiple times in a single read should have an increased weight, because it represents a larger fraction of the read's length. The combination of these terms represents the relative importance of a *k*-mer, and in natural language processing this is known as a *tf-idf* weight (term frequency, inverse document frequency).

Application of *tf-idf* weighting to MinHash sketches is straightforward (Chum et al. 2008). Applied to the read overlapping problem, the weighting is a multiplicative combination of the number of occurrences of a *k*-mer inside a read (the document) and the overall rarity of the *k*-mer among all reads (the corpus). For document similarity, the intuition is that a rare word that occurs multiple times in a single document is a good candidate to identify similar documents. For read overlapping, this statistic has the desirable property that repetitive *k*-mers receive low weights. By reducing the occurrence of repetitive *k*-mers within sketches, the frequency distribution of indexed *k*-mers becomes more uniform. This reduces the number of uninformative, repetitive overlaps that are identified during sketch comparison, significantly improving both runtime and memory usage. Importantly, this is achieved via a probabilistic process so no repeat masking is required and true overlaps between repetitive reads will still be recovered. Alternative weighting schemes are also possible with this technique (e.g. to increase the probability of selecting haplotype-specific *k*-mers), but we focus our evaluation on the *tf-idf* statistic.

We evaluated *tf-idf* weighting on a *Bacillus anthracis* genome sequenced with the Oxford Nanopore MinION (Supplementary Note 1-2). The *B. anthracis* Sterne strain makes a useful test because it possesses a single plasmid often present in multiple copies relative to the main chromosome. In this case, the pXO1 plasmid presented at approximately 6-fold higher coverage than the chromosome (487× vs. 76×). This variable sequencing depth challenges traditional *k*-mer filtering strategies based on a fixed, all-or-nothing threshold. Additionally, it is critically important to recover such plasmids during sequencing, because increased copy number has been previously associated with virulence in other species like *Yersinia pestis* (Wang et al. 2016). As expected, MHAP overlap sensitivity for the plasmid is low (26%) when repetitive *k*-mers are filtered via a fixed threshold. Similarly low sensitivity is seen from Minimap (Li 2016) and DALIGNER (Myers 2014)—17% and 60%, respectively—which both employ a *k*-mer count threshold by default (Supplementary Table S1). Manually increasing this threshold to include plasmid *k*-mers improves Minimap and DALIGNER sensitivity to 94% and 76%, respectively. However, Minimap suffers a drop in positive predictive value (PPV), reporting more false, repeat-induced overlaps. DALIGNER performs a dynamic programing check to confirm all candidate overlaps, so its PPV remains high, but it suffers both a memory (1.6-fold) and runtime (2-fold) penalty. In contrast, Canu's adaptive *tf-idf* weighting scheme requires no parameter adjustment and achieves 89% sensitivity and maintains high PPV (99.5%) with no added runtime or memory penalty.

### Best overlap graph

Canu uses a variant of the greedy “best overlap graph” (BOG) algorithm from (Miller et al. 2008) for constructing a sparse overlap graph. Loading the full overlap graph into memory, as required by string graph formulations (Myers 2005), can be costly for large, complex genomes. In contrast, the greedy algorithm loads only the “best” (longest) overlaps for each read end into memory. This greedy approach is optimal when the read length is sufficiently long (Bresler et al. 2013), and a best overlap graph can be built using just 64 GB of memory for a mammalian genome. However, the greedy algorithm can be misled by repeats that are longer than the overlap length and is therefore prone to mis-assemblies. Canu's new “Bogart” algorithm addresses this problem by statistically filtering repeat-induced overlaps and retrospectively inspecting the graph for potential errors.

In the original BOG method, the best overlaps were selected from a pool of all overlaps below a user-specified error rate threshold, where the overlap error rate is defined as the edit distance divided by the length of the overlap alignment. Thus, this threshold must be set low enough that repeats do not result in false overlaps, yet high enough to account for sequencing error and detect true overlaps. In the new Bogart method, the optimal overlap error rate parameter is automatically estimated from the data, both globally and locally. However, this presents a challenge for raw single-molecule data, which has a sequencing error rate between 10–20% that blurs the distinction between noise and repeat-induced overlaps. Therefore, Canu performs multiple rounds of read and overlap error correction prior to graph construction. After these corrections, the residual read error is estimated from the distribution of all longest overlaps. This full overlap set is then filtered to include only those overlaps within some tolerance of the global median error rate (Figure 2a), and the longest overlaps are recomputed using only this subset. Compared to prior versions of BOG that used a 5% default overlap error rate, Bogart will typically discover an overlap error rate below 2% for corrected single-molecule data. This low threshold effectively removes most false overlaps, allowing the greedy method to construct a clean best overlap graph. From this graph, initial contigs are constructed from the maximal non-branching paths.

**Figure 2.**
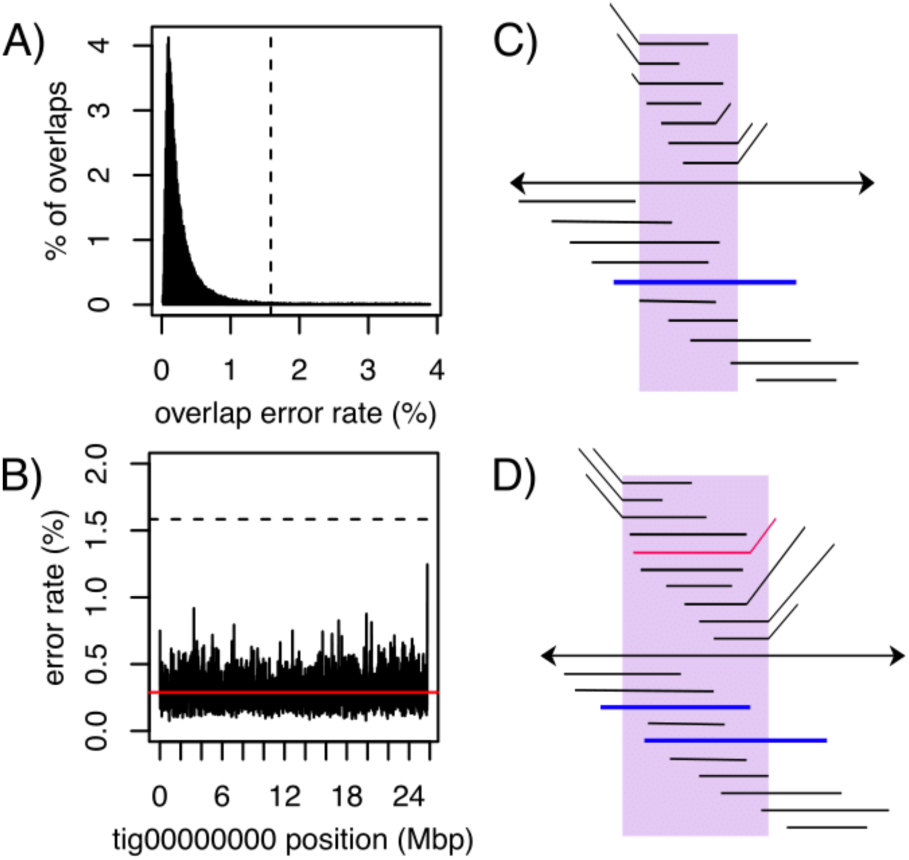
**An illustration of overlap error rate estimation, repeat identification, and splitting**. A) A histogram of all best edge error rates with the auto-selected threshold shown as a dashed line for the *D. melanogaster* PacBio dataset. All overlaps up to 4% error were computed. However, the modal error rate is 0.25% (0.25% median, 0.15% MAD) and Canu chose to use only overlaps below 1.6% error for graph construction on this dataset. B) The dashed line shows the global error rate threshold (1.6%), and the profile shows the locally computed error rate for the largest contig in this assembly. Only overlaps consistent with this local error rate are considered as potential alternate paths when supplementing the initial best overlap graph. By adjusting the error rate for each contig, Canu can separate diverged repeats without making an assumption of uniform read error across the assembly. C) The contig is shown as a black line with arrows on both sides, indicating Bogart extends a path in both the 5′ and 3′ directions until encountering no overlaps or a read that is already incorporated in another contig. Repeat regions annotated by conflicting reads are shown above the contig. The reads align to part of the contig (the repeat) but indicate a different boundary sequence. A single read (blue line) spans the full repeat region, indicating the contig reconstruction is correct. D) Repeat regions annotated by conflicting reads as before. In this case, no single read spans the full repeat region, and the initial contig was built using the overlap between two blue reads. The contig is split if the overlap between the two blue reads is not significantly better than the overlap from either blue read to the conflicting red read.

To evaluate this repeat separation, we compared Canu, Falcon, and Miniasm on a simulated dataset containing two repeat copies with known divergence varying from 0% to 15% and without any spanning reads (Supplementary Note 3). Canu was able to resolve the repeat when the divergence between copies was 3% or higher, without any manual parameter tuning (Supplementary Table S2). In contrast, Falcon could only resolve the repeat at 5%, after it had diverged beyond the default overlap error rate used for corrected reads. Because it lacks a correction step, Miniasm could not resolve the repeat until 13% divergence, higher than the simulated sequencing error rate. This observation may explain why Miniasm is less continuous than Canu and Falcon assemblies on large genomes.

Despite careful correction and overlap filtering, exact or near exact repeats within the error rate tolerance can still add false edges to the graph, resulting in potential mis-assemblies that incorrectly join distant parts of the genome. To guard against this, each initial contig is inspected to identify and correct potential errors. First, the expected overlap error rate for each position of the contig is locally computed using the best overlaps (Figure 2b). Next, all non-best overlaps to reads outside the contig within some deviation of the expected error rate are collected. This excludes sufficiently diverged repeats and haplotypes, while retaining overlaps that are compatible with the local error profile. These overlaps are used to annotate potential alternative branches within the contig and flagged for further inspection. If a branching region is spanned by at least one read (Figure 2c) (Ukkonen 1992) or there is no alternate overlap of similar quality (Figure 2d), it is confirmed as correct. Otherwise, the region is split into at least three new contigs and labeled as an unresolved repeat.

After construction and validation, Canu provides a representation of the final assembly graph in the Graphical Fragment Assembly (GFA) format (Li 2016). This representation is equivalent to a sparse read overlap graph, simplified to remove unambiguous paths and contained reads. Figure 3 shows the Canu assembly graph for *Drosophila melanogaster* sequenced using PacBio. Some chromosome arms are assembled into single contigs, but the graph reveals the structure of the more complex, unresolved repeats in the assembly. For example, chromosome 2L is assembled as a single component in the graph, but is broken towards the end due to a large array of transposable elements and the histone gene cluster, which spans over 500 kbp (Hoskins et al. 2015). These elements also correspond to unfinished gaps in the *D. melanogaster* reference. Canu's graphical output localizes this complex structure to a specific chromosome arm and location. However, the size of the repeats precludes complete assembly. Combining the Canu assembly graph with supplementary long-range information, such as from optical (Hastie et al. 2013) or chromatin contact mapping (Burton et al. 2013; Kaplan and Dekker 2013), could help identify the correct path and resolve such structures.

**Figure 3:**
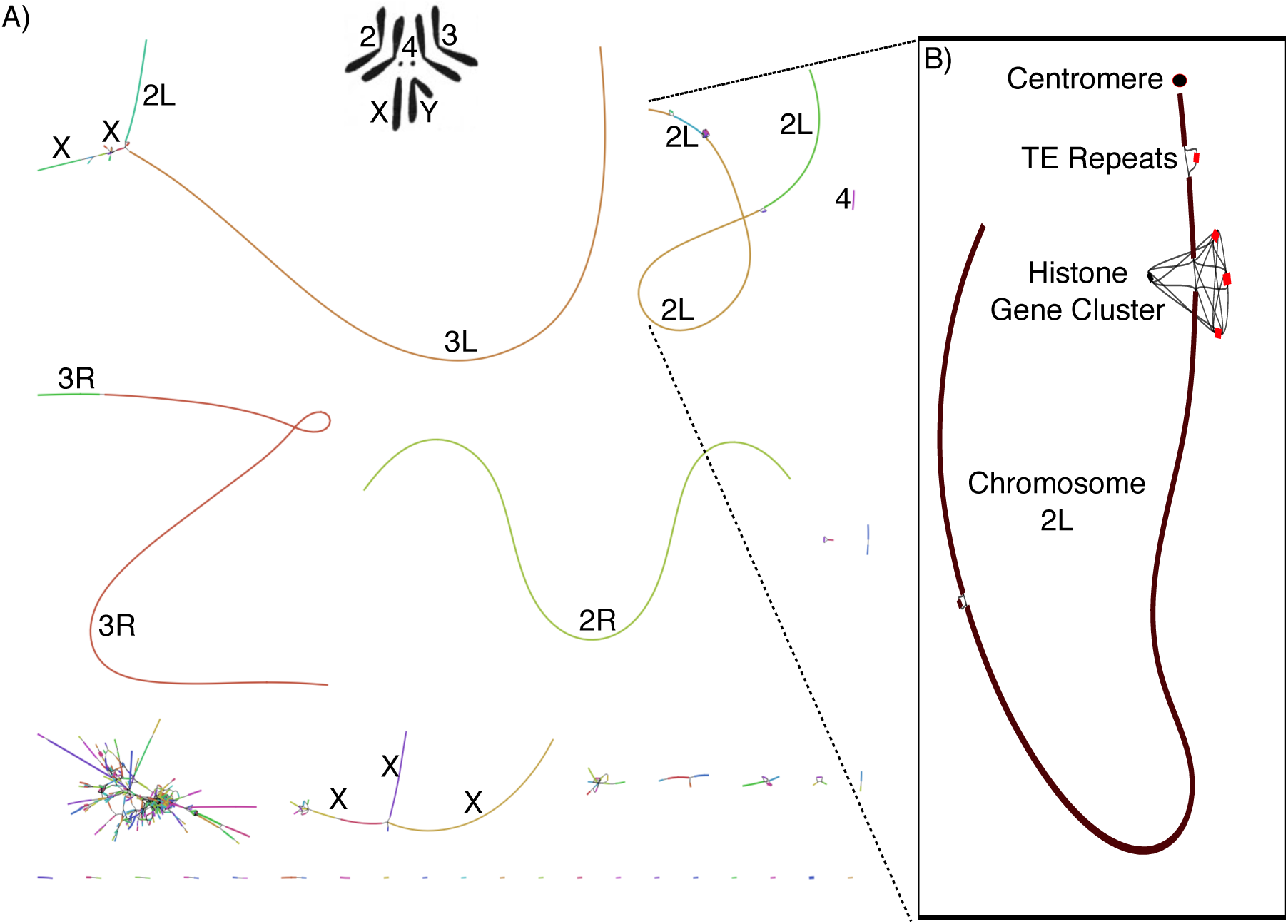
**Canu GFA output localizes complex repeat regions, allowing for improved scaffolding.** A) Bandage (Wick et al. 2015) plot of *D. melanogaster* compared to the karyotype (Stevens 1912; Metz 1914) from FlyBase (Attrill et al. 2016). Nodes are contigs sized by length and edges indicate unused overlaps between contigs. The largest contigs are colored randomly and labeled with their chromosome based on alignment to the reference. B) The callout shows chromosome 2L from positions 3.07 Mbp to 23.12 Mbp, redrawn with the centromere at the top (indicated by a filled circle). Unique contigs are shaded black while repeat contigs are shaded red. While the 2L chromosome scaffold is composed of 10 individual contigs, they are all linked in the output graph. The two red regions correspond to reference gaps at positions 2L:21,485,538, which consists of 100–200 copies of the histone gene cluster spanning over 500 kbp and 2L:22,420,241 which is bordered by several TE repeats (Hoskins et al. 2015). The break in the bottom left of chromosome 2L could not be confidently identified but is next to a feature labeled “FlyBase transposable element” so it is likely a transposable element insertion site. Even though Canu is unable to fully resolve these large repeat arrays, the graph indicates large-scale continuity across chromosome 2L and could enable resolution with secondary technologies.

### Low-coverage hierarchical assembly

Canu substantially lowers the coverage requirements for single-molecule *de novo* assembly. Previously, at least 50× coverage was recommended for hierarchical assembly methods (Berlin et al. 2015; Chakraborty et al. 2016). However, as sequencing lengths and algorithms have improved, so have the minimum input requirements. To quantify performance and determine when a hybrid method may be preferred, we randomly subsampled 10–150× of PacBio P5-C3 coverage from *Arabidopsis thaliana* Ler-0 (Kim et al. 2014) and compared Canu assemblies to both Illumina-only and hybrid assemblies using SPAdes (Antipov et al. 2016). At 20× single-molecule coverage, the Canu assembly is more continuous than the hybrid SPAdes assembly of 20× PacBio combined with 100× Illumina. Although making efficient use of low coverage PacBio data, the hybrid method plateaus after 30×, and the continuity of the Canu 20× assembly is comparable to the best hybrid assembly given 150× of PacBio (Figure 4, Supplementary Note 4, Supplementary Table S3, Supplementary Figure S1). In contrast, Canu continues to improve with increasing PacBio coverage, reaching its maximum assembly continuity around 50×. The amount of improvement is a function of the repeat content and sequence length. PacBio sequence lengths follow a log-normal distribution (Ono et al. 2013), and additional coverage increases the probability of spanning a long repeat. Thus, we would expect continued improvement with higher coverage for larger, more complex genomes. Currently, we recommend the hierarchical method whenever single-molecule coverage exceeds 20×. However, consensus accuracy from low coverage single-molecule data is limited, and polishing (Walker et al. 2014) with short reads is recommended after assembly (Supplementary Table S3).

**Figure 4:**
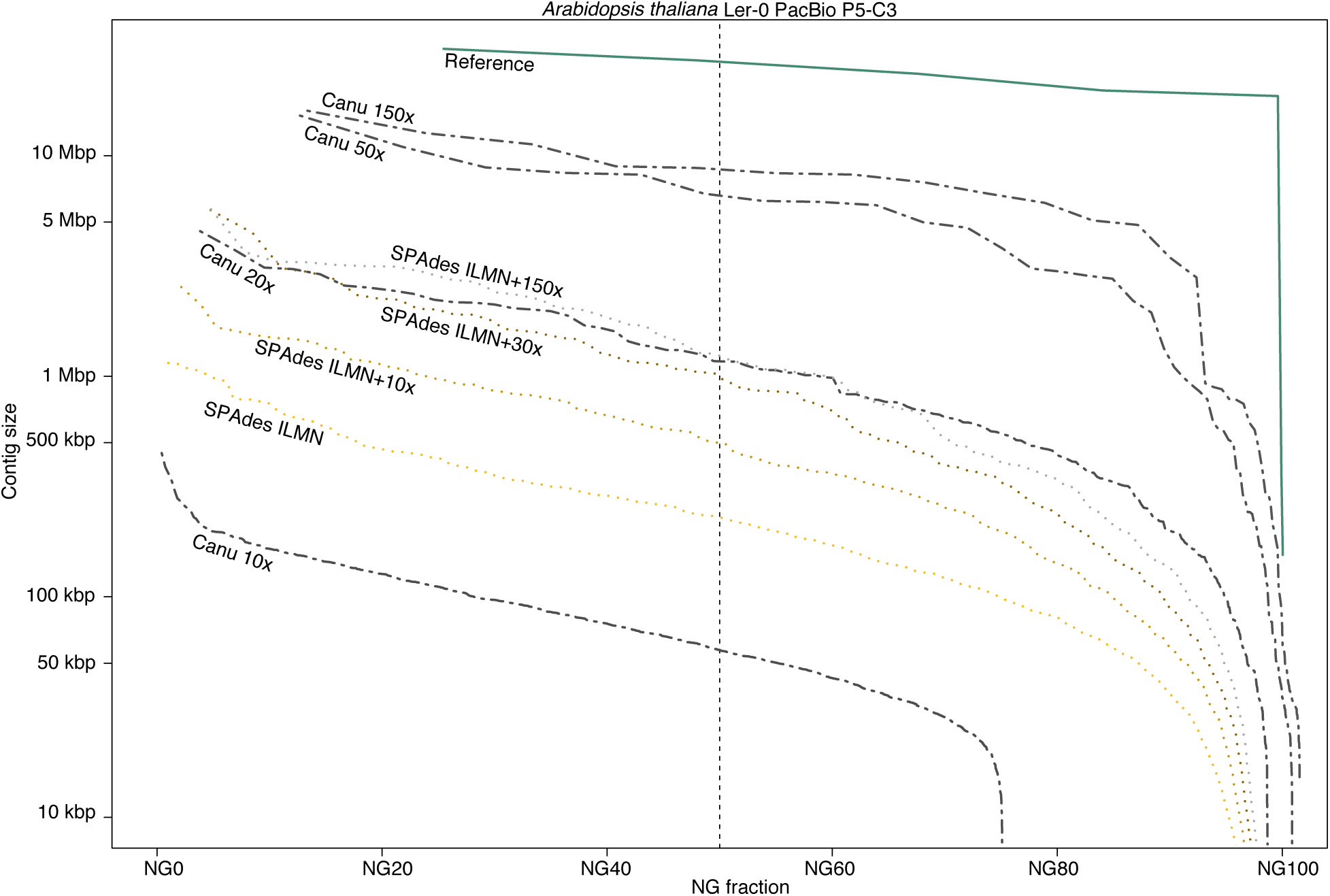
**A comparison of *A. thaliana* assembly continuity for Canu and SPAdes.** Each set of contigs is sorted from longest to shortest and plotted versus a cumulative percentage of the genome covered. Assemblies with larger contigs appear in the top of the plot. The ideal assembly corresponds to the green reference line. The commonly used NG50 metric corresponds to the vertical dashed line. Canu quickly gains continuity with increasing coverage, approaching the limit with 50× PacBio on this genome. In contrast, while making a large gain from Illumina-only to 10× PacBio, SPAdes continuity plateaus by 30×, and the Canu 20× assembly is comparable to the hybrid SPAdes assembly using 150× PacBio.

### Assembly evaluation

We evaluated Canu on a variety of microbial and eukaryotic genomes, and compared with Falcon (Chin et al. 2016), Miniasm (Li 2016), and hybrid SPAdes (Antipov et al. 2016) using both PacBio and Oxford Nanopore sequencing data (Supplementary Note 5–7). Continuity was measured using maximum and NG50 contig size, where NG50 is the longest contig such that contigs of this length or greater sum to at least half the haploid genome size. Accuracy was computed via alignment to the nearest available reference genome using MUMmer (Kurtz et al. 2004), and reported using the GAGE (Salzberg et al. 2012) metrics, which evaluate both base (single nucleotide) and structural breakpoints (inversions, relocations, and translocations). An ideal assembly has high continuity, low breakpoints, and high base accuracy, with 99.99% (Phred QV40 (Ewing and Green 1998)) commonly defined as the minimum quality for a “finished” sequence (Felsenfeld et al. 1999; Schmutz et al. 2004).

### PacBio sequence assembly

We assembled bacterial and eukaryotic genomes recently released (Kim et al. 2014) and available from PacBio DevNet (https://github.com/PacificBiosciences/DevNet/wiki/Datasets). Table 1 shows that Canu produces the most continuous assembly on three of the four eukaryotic genomes tested, while maintaining high accuracy (Supplementary Figure S2–S6). In the one case that Miniasm produces a higher NG50 (*Caenorhabditis elegans*), both Falcon and Miniasm introduce large-scale structural rearrangements not present in the Canu assembly (Supplementary Figure S5). For initial assembly, Miniasm (Li 2016) is an order of magnitude faster than Canu and Falcon (Supplementary Note 8, Supplementary Table S4–S7). However, in contrast to Canu and Falcon, Miniasm does not perform a gapped alignment for either overlapping or consensus generation. Instead, Miniasm generates a string graph (Myers 2005) directly from approximate read overlaps and labels the edges of this graph with the raw read sequences. Thus, the average identity of the resulting assembly is equal to the identity of the input sequences, and the approximate overlap positions can leave large insertions and deletions in the assembly at the boundaries of the read segments. As a result, the Miniasm assemblies have both low base accuracy (<90%) and a higher frequency of large insertions and deletions, which can be difficult to remove during polishing. Therefore, Miniasm requires four rounds of Quiver polishing (Chin et al. 2013) before the assembly quality converges, whereas Canu requires only a single polishing round and is ultimately fastest to generate a polished assembly (Table 1, Supplementary Note 9, Supplementary Table S8–S11). To test if Miniasm polishing could be accelerated using a different algorithm, we tested the recently released Racon tool (Vaser et al. 2016), which was designed for this purpose. However, on *C. elegans*, two rounds of Racon required 60 CPU hours and produced a lower quality consensus than a single round of Quiver, which required a comparable 110 CPU hours.

**Table 1:** Canu is fastest for generating a high-quality polished assembly from PacBio data

| Genome | Asm/Polish | Max (Mbp) | NG50 (Mbp) | % Ref | # Breakpoints | Time (CPU h) | % Idy |
| --- | --- | --- | --- | --- | --- | --- | --- |
| *E. coli* | Canu+Quiver | **4.68** | **4.68** | **100%** | **0** | 12.25 | **99.9999%** |
|  | Falcon+Quiver | 4.64 | 4.64 | **100%** | 2 | 25.14 | 99.9998% |
|  | Miniasm+Quiver | 4.64 | 4.64 | 99.99% | 2 | 31.93 | 99.9998% |
|  | SPAdes | 4.64 | 4.64 | **100%** | **0** | **4.09** | 99.9972% |
| *D. melanogaster* | Canu+Quiver | **25.78** | **21.31** | **97.47%** | 1,025 | **1,396.52** | 99.9795% |
|  | Falcon+Quiver | 23.08 | 9.84 | 96.12% | 1,054 | 2,305.92 | **99.9813%** |
|  | Miniasm+Quiver | 15.85 | 5.84 | 96.51% | **752** | 1,484.33 | **99.9813%** |
| *A. thaliana* | Canu+Quiver | **15.95** | **8.31** | **82.94%** | 220 | **925.31** | **99.0710%** |
|  | Falcon+Quiver | 15.94 | 8.17 | 82.72% | 222 | 1,132.25 | **99.0710%** |
|  | Miniasm+Quiver | 11.61 | 5.07 | 82.88% | **205** | 976.43 | **99.0710%** |
| *C. elegans* | Canu+Quiver | 5.34 | 2.35 | **99.70%** | 139 | 410.07 | **99.9745%** |
|  | Falcon+Quiver | 4.99 | 1.88 | 98.82% | **138** | **397.40** | 99.9735% |
|  | Miniasm+Quiver | **5.85** | **2.96** | 99.44% | 141 | 526.16 | 99.9706% |
| CHM1 | Canu+Quiver | **80.08** | **21.95** | **86.84%** | 1,105 | **22,749.71** | 99.8081% |
|  | Falcon+Quiver | 52.34 | 9.46 | 86.58% | **1,082** | 68,789.00 | **99.8086%** |

Canu shows good scaling to mammalian genomes, completing a polished human assembly tenfold faster than Celera Assembler 8.2, which was used to assemble the first human genome from PacBio data alone (Berlin et al. 2015), and threefold faster than the more recent Falcon assembler (Supplementary Table S4–S5). Canu runtime improvements come from recent optimizations to the initial overlapping and read correction process (Methods), which have traditionally been the slowest step in hierarchical assembly. Read correction is now the fastest step of the Canu pipeline. As a result, Canu is often able to generate a complete assembly in less time than Falcon requires for its initial DALIGNER (Myers 2014) overlapping stage (Supplementary Table S4–S5). On the human genome, where the upfront cost of building MHAP sketches is most effectively amortized, Canu's initial overlapping step is also faster than Minimap (Supplementary Table S6), but Miniasm failed to assemble this dataset due to its in-memory string graph construction, which exceeded 1 TB of memory. Canu's greedy algorithm required less than 36 GB for the same dataset.

Canu also represents a dramatic improvement over the latest version of Celera Assembler (Berlin et al. 2015). Our previous PacBio P5-C3 human (CHM1) assembly required >250,000 CPU hours with Celera Assembler, resulting in a contig NG50 of 4 Mbp (Berlin et al. 2015). The re-assembly of this same dataset with Canu required <25,000 CPU hours and the NG50 increased to over 7 Mbp. Improvements to PacBio chemistries are also resulting in impressive assembly gains. An updated assembly using the more recent PacBio P6-C4 chemistry requires the same runtime, yet increases the NG50 5-fold to over 20 Mbp. This *de novo* Canu assembly has comparable assembly size, contig counts, and continuity to the human reference assemblies before NCBI Build 34 (ca. 2003), which is the release immediately prior to the “finished” human genome (International Human Genome Sequencing 2004). The contig sizes of this Canu human assembly are also comparable to the scaffold sizes generated by Celera (Istrail et al. 2004), which used Sanger sequencing with a range of insert sizes and BACs.

Since CHM1 is effectively a haploid sample, we also tested Canu on the recently released diploid Chinese human genome (HX1) (Shi et al. 2016). This data has a similar read length distribution to the CHM1 P5-C3 data (Supplementary Figure S7), albeit at twice the coverage, so one would expect a slight continuity improvement. As expected, the Canu HX1 assembly achieved an NG50 of 9.00 Mbp (Supplementary Figure S8), improving on the published Falcon assembly NG50 of 7.61 Mbp (Shi et al. 2016) and thereby demonstrating that Canu performs equally well on diploid human genomes. However, due to the relatively low level of heterozygosity, Canu will currently collapse human haplotypes and would require dedicated phasing to generate a haplotype-resolved human assembly.

### Nanopore sequence assembly

Currently, the Oxford Nanopore MinION can read either one or both strands of a double-stranded DNA molecule. The “1D” mode sequences only the template strand, whereas the “2D” mode sequences both the template and complement strands via a hairpin adapter. This technique is similar to PacBio circular consensus sequencing (CCS) (Travers et al. 2010). Because the 2D mode provides two independent observations of each base, the per-read accuracy is improved (e.g. from 70% to 86% for R7.3 chemistry (Figure 5a)). To date, all assembly evaluations have focused on the more accurate 2D sequences (Loman et al. 2015; Judge et al. 2016; Sovic et al. 2016). While more accurate, the library preparation for 2D sequencing is more complex, reduces the effective throughput of the instrument (each molecule must be read twice), and currently produces shorter sequences. Thus, we designed Canu to assemble both 2D and the noisier 1D sequences, which benefit from increased read length and throughput, both key factors for genome assembly.

**Figure 5:**
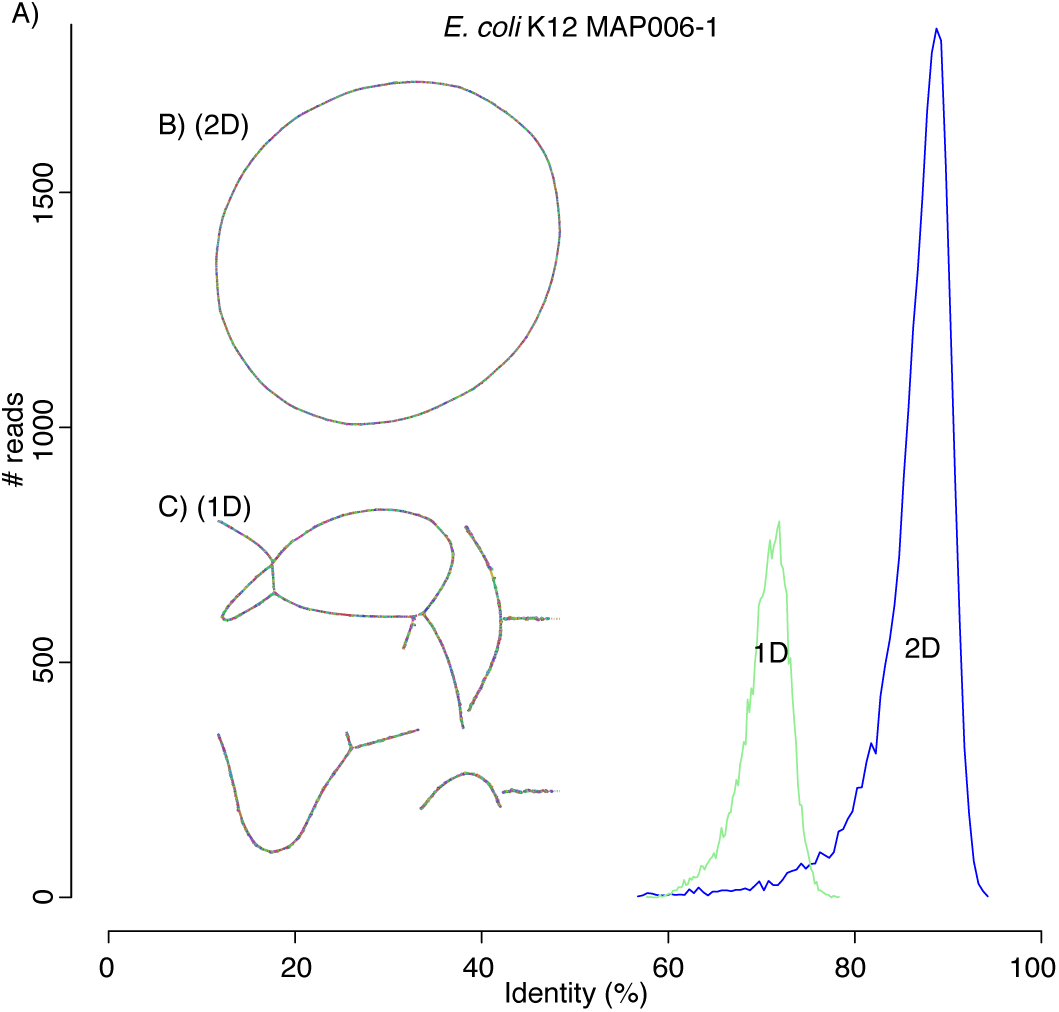
**Canu can assemble both 1D and 2D Nanopore *E. coli* reads.** A) A comparison of error rates for 1D and 2D read error rates versus the reference. Template 1D and 2D reads from the MAP006-1 *E. coli* dataset were aligned independently to compute an identity for all reads with an alignment over 90% of their length (95% of the 2D reads and 86% of the 1D reads had an alignment over 90% of their length). The 2D sequences averaged 86% identity and the 1D reads averaged 70% identity. B) Bandage plot of the Canu best overlap graph for the 2D data. The genome is in a single circle representing the full chromosome. C) The corresponding plot for 1D data. While highly continuous, there are multiple components due to missed overlaps and unresolved repeats (due to the higher sequencing error rate).

Table 2 shows Canu assemblies of seven recent 2D Nanopore sequencing runs: http://lab.loman.net/2015/09/24/first-sqk-map-006-experiment/and http://lab.loman.net/2016/07/30/nanopore-r9-data-release/ (Loman et al. 2015). Consistent with independent evaluations (Judge et al. 2016; Sovic et al. 2016), Canu produces highly continuous assemblies from Nanopore data alone, and the continuity of Canu assemblies was equal to or better than all assemblers tested. Miniasm was again extremely fast and produced structurally correct and continuous assemblies (Supplementary Note 10, Supplementary Table S12–S14, Supplementary Figure S9–S15), except for *B. anthracis*, where it failed to assemble the high-copy plasmid pXO1 due to its stringent *k*-mer filtering. As with PacBio, the initial Minimap assemblies also have low base accuracy. For Nanopore data, Minimap assemblies were less than 90% accurate, whereas Canu assemblies typically exceeded 99%. Consensus polishing using the Nanopore signal data with Nanopolish (Loman et al. 2015) further improved the accuracy of all assemblies to as high as 99.85%, but polishing the lower quality Miniasm assemblies to comparable accuracy was 750% slower (Supplementary Table S12–S14).

**Table 2:** Canu consistently assembles complete genomes from only Oxford Nanopore data

| Genome | Asm/Polish | # Contigs | Max (Mbp) | % Ref | # Breakpoints | Time (CPU h) | % Idy |
| --- | --- | --- | --- | --- | --- | --- | --- |
| *E. coli* MAP005 | Canu+Nanopolish | (1) | **4.64** | **99.98%** | 2 | 376.87 | **99.43%** |
|  | Falcon+Nanopolish | 105 | 0.42 | 23% | 2 | **106.2** | 99.41% |
|  | Miniasm+Nanopolish | 3 | 3.40 | 99.96% | **0** | 2,344.02 | 99.36% |
| *E. coli* MAP006-1 | Canu+Nanopolish | (1) | 4.63 | 99.80% | **0** | **167.04** | **99.81%** |
|  | Falcon+Nanopolish | (1) | 4.63 | 99.86% | **0** | 207.45 | 99.78% |
|  | Miniasm+Nanopolish | (1) | **4.66** | **99.97%** | 2 | 1,801.02 | 99.72% |
| *E. coli* MAP006-2 | Canu+Nanopolish | (1) | 4.64 | 99.91% | **2** | **168.69** | **99.78%** |
|  | Falcon+Nanopolish | (1) | 4.64 | **99.94%** | **2** | 196.16 | 99.76% |
|  | Miniasm+Nanopolish | (1) | **4.65** | 99.70% | 4 | 1,482.95 | 99.69% |
| *E. coli* MAP006-PCR-1 | Canu+Nanopolish | (1) | **4.64** | 99.95% | **0** | **164.08** | **99.84%** |
|  | Falcon+Nanopolish | (1) | 4.63 | 99.80% | 2 | 168.37 | 99.82% |
|  | Miniasm+Nanopolish | 3 | 2.15 | **99.96%** | **0** | 1,338.28 | 99.77% |
| *E. coli* MAP006-PCR-2 | Canu+Nanopolish | (1) | 4.64 | 99.99% | 2 | **206.09** | **99.85%** |
|  | Falcon+Nanopolish | (1) | 4.64 | **100.00%** | 2 | 212.89 | 99.84% |
|  | Miniasm+Nanopolish | (1) | **4.65** | 99.98% | **0** | 1,669.83 | 99.81% |
| *B. anthracis* | Canu+Nanopolish | (2) | 5.20 | **99.77%** | **0** | 894.40 | 99.14% |
|  | Falcon+Nanopolish | 31 | 0.47 | 86.29% | **0** | **795.93** | **99.17%** |
|  | Miniasm+Nanopolish | 4 | **5.22** | 97.21% | **0** | 5,094.90 | 99.05% |
| *Y. pestis* | Canu+Nanopolish | (4) | 4.67 | **99.97%** | **11** | **254.25** | **99.76%** |
|  | Falcon+Nanopolish | (4) | **4.68** | **99.97%** | 12 | 295.01 | 99.72% |
|  | Miniasm+Nanopolish | 9 | 2.69 | 99.91% | **11** | 2,000.16 | 99.65% |

Generating a finished-quality (>99.99%) consensus sequence from Nanopore reads required polishing with complementary short-read data. We repeated the above evaluation, but substituted Pilon (Walker et al. 2014) for Nanopolish (Loman et al. 2015), and included comparisons to hybrid SPAdes (Table 3). Pilon aligns Illumina reads against an assembled sequence and corrects base errors and small insertions and deletions (Indels). As with Nanopolish, this process was iterated until consensus quality converged, except for hybrid SPAdes, which did not require additional polishing. Combined assembly and polishing times for all assemblers were comparable. Canu, Falcon, and SPAdes routinely exceeded 99.99% polished base accuracy, but Miniasm was unable to exceeded 99.9% after many rounds of polishing (Supplementary Table S15). The residual Miniasm errors were large (average >500 bp) expansions or collapses in the draft assembly (Supplementary Figure S16), which are difficult to correct using short-read sequences. Hybrid SPAdes was typically most accurate, both in terms of base and structural accuracy. However, on the repetitive *Y. pestis* genome, it was significantly less continuous than hierarchical methods, and on the newer high-quality Nanopore datasets, the polished Canu accuracy exceeded SPAdes (Supplementary Note 11, Supplementary Table S16–S19, Supplementary Figure S17–S23).

**Table 3.** Nanopore assemblies exceed hybrid methods in continuity and match their quality when polished with Illumina data

| Genome | Asm/Polish | # Contigs | Max (Mbp) | % Ref | # Breakpoints | Time (CPU h) | % Idy |
| --- | --- | --- | --- | --- | --- | --- | --- |
| *E. coli* MAP005 | Canu+Pilon | (1) | **4.65** | 99.99% | 2 | 10.98 | 99.9873% |
|  | Falcon+Pilon | 105 | 0.42 | 23.04% | 2 | 4.36 | 99.9550% |
|  | Miniasm+Pilon | 3 | 3.40 | 90.62% | 42 | **3.15** | 97.3878% |
|  | SPAdes | (1) | 4.64 | **100.00%** | **0** | 3.61 | **99.9989%** |
| *E. coli* MAP006-1 | Canu+Pilon | (1) | 4.63 | 99.82% | **0** | 5.89 | **99.9995%** |
|  | Falcon+Pilon | (1) | 4.63 | 99.86% | **0** | 7.3 | 99.9964% |
|  | Miniasm+Pilon | (1) | **4.66** | 96.97% | 21 | **3.14** | 99.6118% |
|  | SPAdes | (1) | 4.64 | **100.00%** | **0** | 3.65 | 99.9965% |
| *E. coli* MAP006-2 | Canu+Pilon | (1) | **4.64** | 99.94% | 2 | 3.92 | **99.9987%** |
|  | Falcon+Pilon | (1) | **4.64** | 99.94% | 2 | 3.93 | 99.9933% |
|  | Miniasm+Pilon | (1) | **4.64** | 97.98% | 26 | **2.73** | 99.6336% |
|  | SPAdes | (1) | **4.64** | **100.0%** | **0** | 3.56 | 99.9965% |
| *E. coli* MAP006-PCR-1 | Canu+Pilon | (1) | **4.64** | 99.95% | **0** | 4.15 | **99.9993%** |
|  | Falcon+Pilon | (1) | 4.63 | 99.80% | 2 | 3.55 | 99.9969% |
|  | Miniasm+Pilon | 3 | 2.16 | 98.41% | 12 | **2.15** | 99.6734% |
|  | SPAdes | 2 | 3.95 | **100.00%** | **0** | 3.56 | 99.9965% |
| *E. coli* MAP006-PCR-2 | Canu+Pilon | (1) | 4.64 | **100.00%** | 2 | 6.16 | **99.9992%** |
|  | Falcon+Pilon | (1) | 4.64 | **100.00%** | 2 | 9.22 | 99.9963% |
|  | Miniasm+Pilon | (1) | **4.65** | 98.57% | 20 | **2.69** | 99.6734% |
|  | SPAdes | (1) | 4.64 | **100.00%** | **0** | 4.00 | 99.9965% |
| *B. anthracis* | Canu+Pilon | (2) | 5.21 | 99.77% | 1 | 65.01 | 99.8476% |
|  | Falcon+Pilon | 31 | 0.48 | 86.31% | **0** | 14.95 | 99.8888% |
|  | Miniasm+Pilon | 4 | **5.25** | 79.36% | 44 | **4.9** | 92.2732% |
|  | SPAdes | 6 | 4.13 | **100.00%** | **0** | 8.47 | **99.9948%** |
| *Y. pestis* | Canu+Pilon | (4) | **4.66** | **99.83%** | 23 | 17.92 | 99.8946% |
|  | Falcon+Pilon | (4) | 4.64 | 99.65% | 26 | 10.63 | 99.8715% |
|  | Miniasm+Pilon | 9 | 2.70 | 93.76% | 42 | **8.68** | 98.7866% |
|  | SPAdes | 29 | 0.37 | 95.99% | **15** | 17.08 | **99.9559%** |

### Nanopore 1D sequence assembly

We evaluated the performance of Canu on noisy 1D data using only the template sequences from the *Escherichia coli* MAP006-1 dataset, which averaged a raw 1D accuracy of just 70% (Figure 5a). To deal with this high error, we exploited the modularity of Canu to run ten rounds of correction, with the output of each round fed as input to the next (Supplementary Note 12). The corrected reads were then assembled into ten contigs with an NG50 of 619 kbp and a maximum contig size of 1.22 Mbp covering 89% of the reference at 85.52% identity versus a single circular chromosome for 2D data (Figure 5b-c). In contrast, the Miniasm assembly of this data covered less than 10% of the reference at 76.76% identity (Supplementary Figure S24). Polishing the Canu assembly with Nanopolish converged on a 1D consensus accuracy of 98% identity, and short-read polishing with Pilon improved the assembly to 93.83% coverage and 99.72% identity. Thus, despite their high error, we conclude that 1D sequences as low as 70% identity can be assembled, albeit at reduced consensus quality. However, more recent Nanopore sequencing chemistries are producing 1D reads with 85% accuracy, for which only a single round of correction is necessary.

Few eukaryotic Nanopore datasets are currently available due to the low throughput of the initial MinION instruments. However, as previously demonstrated using PacBio data, Canu easily scales to mammalian-sized genomes, and as Nanopore throughput improves it is expected that highly continuous eukaryotic assemblies will be possible. For an early test, we assembled the *Saccharomyces cerevisiae* genome from available R6 and R7 MinION data (Goodwin et al. 2015). This older dataset contains only 20× coverage of 2D reads and an average identity of 70% (Figure 6a), significantly lower than produced by newer chemistries. Despite this, Canu was able to assemble the dataset using the same iterative correction strategy as for 1D reads (Figure 6b, Supplementary Note 13, Supplementary Figure S25). The resulting assembly comprises 41 contigs, with a majority of chromosomes in one or two contigs and an NG50 of 469 kbp covering 95.22% of the reference at 94.33% identity. Illumina polishing with Pilon improved the assembly to 96.86% coverage at 99.83% identity. Prior to Canu, this dataset could only be assembled via a hybrid approach. Newer Nanopore chemistries are not expected to require an iterative correction strategy, and improved instrument throughput will enable fully assembled yeast chromosomes (Istace et al. 2016).

**Figure 6:**
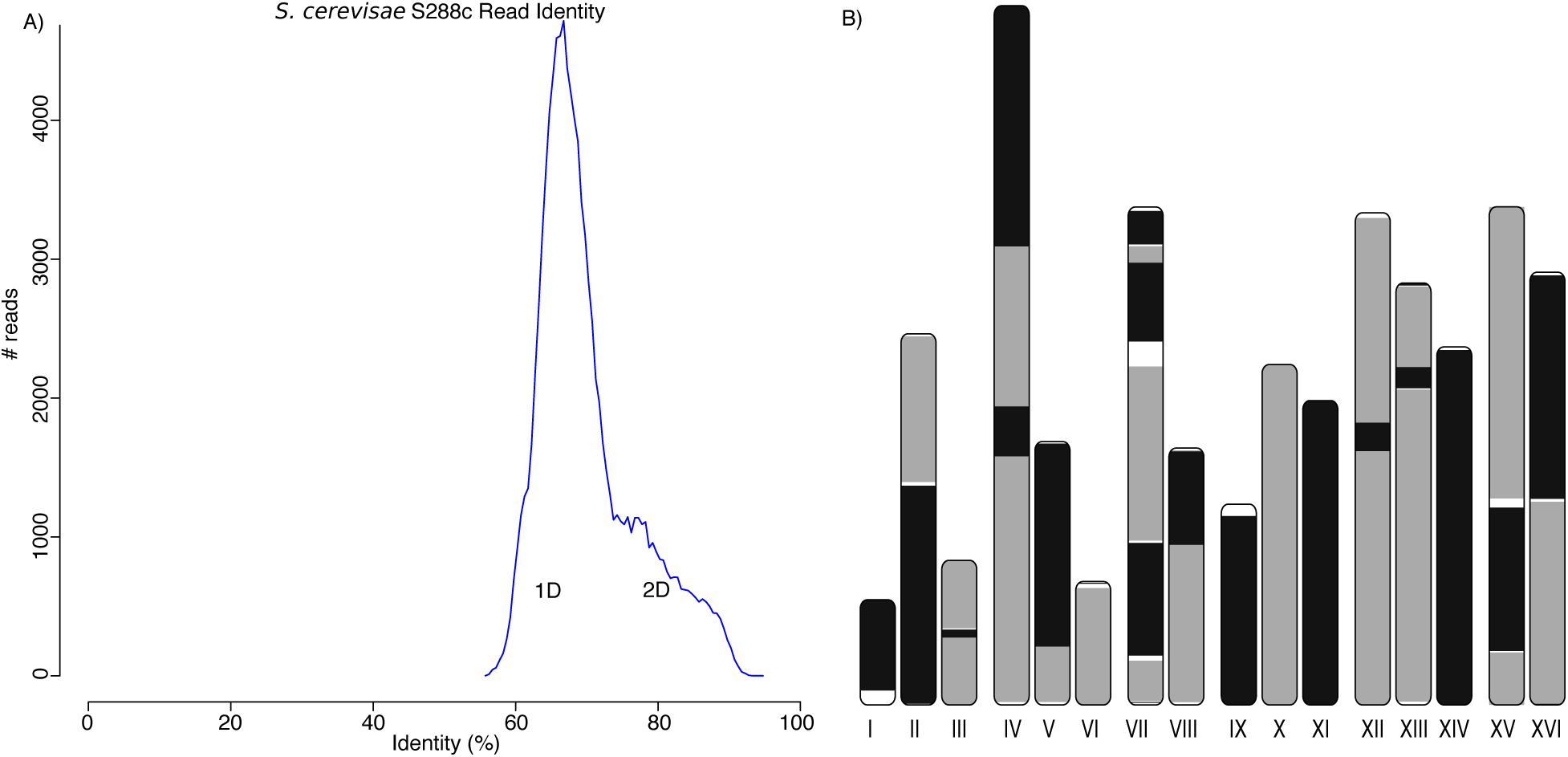
**A highly continuous *S. cerevisae* assembly from noisy 1D and 2D MinION reads.** A) A histogram of read error rates (1D and 2D) versus the reference. Alignment identity was computed only for reads with an alignment over 90% of their length. The majority of reads were below 75% identity with an overall average of 70%. B) Assembled Canu contigs were aligned to the reference and all alignments over 1 kbp in length and >90% identity were then plotted using the coloredChromosomes package (Böhringer et al. 2002). Alternating shades indicate adjacent alignments, so each transition from gray to black represents a contig boundary or alignment breakpoint. White regions indicate regions missing from the assembly. Most chromosomes are in less than 3 contigs, indicating structural agreement with the reference.

## Discussion

Canu is able to generate highly continuous assemblies from both PacBio and Nanopore sequencing, but signal-level polishing is required to maximize the final consensus accuracy. Such algorithms use statistical models of the sequencing process to predict base calls directly from the raw instrument data, which is a richer source of information than FASTQ Phred quality values. Currently, a PacBio base accuracy of 99.999% (QV50) is achievable with Quiver polishing (Chin et al. 2013; Koren et al. 2013), but Nanopore is limited to at most 99.9% (QV30) with Nanopolish (Loman et al. 2015) due to systematic sequencing errors (Goodwin et al. 2015). Both tools are technology specific and must be trained on each new chemistry, so future improvements are possible. Alternatively, complementary short-read sequencing can be used for consensus polishing with Pilon. On recent Nanopore sequencing data, Illumina-polished Canu assemblies can reach QV50 and exceed the base accuracy of hybrid SPAdes assemblies. Thus, the combination of Nanopore and Illumina sequencing provides a new alternative for the generation of finished microbial genomes. However, due to the difficulty of mapping short Illumina reads to repeats, signal-polished PacBio assemblies currently deliver the highest overall quality.

Canu assembly followed by either single-molecule or short-read polishing is an efficient method for generating high-quality assemblies. Our results indicate that while Miniasm (Li 2016) can rapidly produce continuous and structurally accurate assemblies, the multiple rounds of polishing needed to produce an accurate consensus sequence becomes a computational bottleneck. Additionally, Canu is the only tool capable of assembling low-accuracy 1D Nanopore data, while scaling to gigabase-sized genomes—an important application given the pending release of high-throughput Nanopore sequencers. Combined with Canu's adaptive *k*-mer weighting strategy, the assembly of repetitive heterochromatic sequence may be possible with high-coverage, long-read nanopore sequencing.

Canu currently splits haplotypes into separate contigs wherever the allelic divergence is greater than the post-correction overlap error rate. This threshold is typically 1.5% for recent PacBio data. This splitting results in an assembly size larger than the haploid genome size. Although these regions are kept separate in the assembly graph, no effort is currently made to annotate such regions or phase multiple bubbles into larger haplotype blocks. Less diverged haplotypes, such as human, are collapsed, as demonstrated by the HX1 dataset. Currently, only abundance is considered for *k*-mer weighting, which avoids the consideration of false, repetitive overlaps. However, this same scheme could be used to improve the discrimination of minor variants between repeats and haplotypes by preferring haplotype-specific *k*-mers during sketch construction. This would increase the power of Canu's statistical overlap filter, which prevents the merging of diverged repeats and haplotypes.

For further improved haplotype reconstruction, it would be possible to apply an approach like Falcon-Unzip (Chin et al. 2016) to the Canu assembly graph to generate phased contigs based on linked variants identified within the single-molecule reads. For repeat structures, the current algorithm can resolve any repeat copy with more divergence than the post-correction overlap error rate. In the future, similar repeats could be resolved using more sophisticated graph traversals. For example, if one copy of a two-copy repeat is spanned, a correct reconstruction of the unspanned copy can be inferrered given that the other copy is correctly assembled (Ukkonen 1992). Alternatively, secondary information from technologies like 10× Genomics (Zheng et al. 2016) or Hi-C (Selvaraj et al. 2013) could be used to guide walks through the Canu graph. Ultimately, because Hi-C provides megabase-scale linkage information, the integration of this technology with Canu assembly graphs could lead to complete *de novo* assemblies that span entire mammalian chromosomes from telomere to telomere, as was recently demonstrated for the domestic goat genome (Bickhart et al. 2016).

## Methods

### Architecture

Canu is a modular assembly infrastructure comprised of three primary stages—correction, trimming, and assembly (Figure 1)—that can be run on a single computer or multi-node compute cluster. For multi-node runs, recommended for large genomes, Canu supports Sun Grid Engine (SGE), Simple Linux Utility for Resource Management (SLURM), Load Sharing Facility (LSF), and Portable Batch System (PBS)/Torque job schedulers. Users without access to an institutional compute cluster can run large Canu assemblies via a cloud-computing provider using toolkits such as StarCluster (http://star.mit.edu/cluster/).

As a Canu job progresses, summary statistics are updated in a set of plaintext and HTML reports. The primary data interchange between stages is FASTA or FASTQ inputs, but for efficiency each stage stores input reads in an indexed database, after which the original input is no longer needed. Each of the three stages begins by identifying overlaps between all pairs of input reads. Although the overlapping strategy varies for each stage, each counts *k*-mers in the reads, finds overlaps between the reads, and creates an indexed store of those overlaps. By default the correction stage uses MHAP (Berlin et al. 2015) and the remaining stages use overlapInCore (Myers et al. 2000). From the input reads, the correction stage generates corrected reads; the trimming stage trims unsupported bases and detects hairpin adapters, chimeric sequences, and other anomalies; and the assembly stage constructs an assembly graph and contigs. The individual stages can be run independently or in series.

For distributed jobs, local compute resources are polled to build a list of available hosts and their specifications. Next, based on the estimated genome size, Canu will choose an appropriate range of parameters for each algorithm (e.g. number of compute threads to use for computing overlaps). Finally, Canu will automatically choose specific parameters from each allowed range so that usage of available resources is maximized. As an example, for a mammalian sized genome, Canu will choose between 1 to 8 compute threads and 4 to 16 GB memory for each overlapping job. On a grid with ten hosts, each with 18 cores and 32 GB of memory, Canu will maximize usage of all 180 cores by selecting 6 threads and 10 GB of memory per job. This process is repeated for each step, and allows automated deployment across varied cluster and host configurations, simplifying usage and maximizing resource utilization.

### MinHash Overlapping

Canu uses an updated version of the MinHash Alignment Process (MHAP) for computing all-versus-all overlaps from noisy, single-molecule sequences (Berlin et al. 2015). MHAP has been further optimized for both speed and accuracy since the initial version. As described below, the most substantial algorithmic changes involve the sketching and filtering strategies. MHAP uses a two-stage overlap filter, where the first stage identifies read pairs that are likely to share an overlap and the second stage estimates the extent and quality of the overlap. For the first stage, MHAP now implements *tf-idf* weighting to prefer informative, non-repetitive *k*-mers. This increases sensitivity to true overlaps, while reducing the number of false, repetitive overlaps considered. For the second stage, MHAP now implements a “bottom sketch” strategy similar to Mash (Ondov et al. 2016), which significantly decreases memory usage and runtime. The Mash distance formula is also used to estimate the error rate (quality) of the identified overlaps directly from the sketches, without the need for a gapped alignment (Ondov et al. 2016). Engineering improvements include a switch to the FastUtil (http://fastutil.di.unimi.it) hash table implementation, which resulted in a 3-fold speedup, and an increase in the maximum *k*-mer size from 16 to 128 to support greater specificity on low-error datasets. Overall, the new MHAP version is 10-fold faster, on average, and over 40-fold faster on mammalian genomes that the original version, while maintaining similar accuracy.

There have been several *tf-idf* formulations proposed for document and image retrieval (Manning et al. 2008), but for our purposes we use:

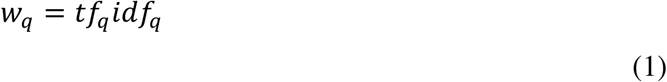

For each read, *tf_q_* is the number of occurrences of *k*-mer *q* in the read, and *idf_q_* observed across all reads. Specifically, for all *k*-mers in the input read set, let *f_max_* be the maximum observed frequency, *f_min_* be the minimum observed frequency, and *f_q_* be the frequency of a specific *k-mer q*. By default, only 0.0005% of the most abundant *k*-mers are recorded, and all others are assigned *f_min_*. We define *idf_q_* as:

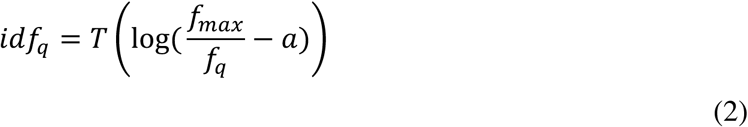

The parameter *a* ∈ [0,1] controls how strongly less common *k*-mers are preferred in relation to the more common ones, and *T* linearly transforms the values between 1 and *idf_max_*, the maximum allowed weight. The minimum possible value is computed by plugging the maximum observed frequency of the most popular *k*-mer into formula (2), and the maximum possible value is computed by plugging in the filter cutoff value provided to MHAP (5×10^-6^ being the default). The idf values are then linearly rescaled to fall in the range [1,*idf_max_*]. Any *k*-mer that does not exist in the filter file is assigned *idf_max_*.

For a general positive floating point number, (Chum et al. 2008) provided a formula for directly computing the *w*-weighted hash value for MinHash ranking. However, this formula requires computing *s · L* logarithms to generate a sketch, which is computationally expensive (where *s* is the sketch size and *L* is the read length). As in the original MHAP implementation, we compute the initial hash value using the MurmurHash3 hash (http://code.google.com/p/smhasher/wiki/MurmurHash3), while the subsequent *s* – 1 hashes are computed from a pseudorandom number generator (Berlin et al. 2015). We discretize the *tf-idf* to a limited range using rounding, which requires at most (*s* – 1) · *L* · *w_max_* random number computations, where *w_max_* is the maximum weight computed by MHAP, which is comparatively faster. We use *idf_max_* = 3 and *a* = 0.9 by default as a compromise between speed and performance.

Recall that MinHash selects which *k*-mers will be included in the sketch on the basis of their hash value. In the original MHAP implementation, a set *Γ* of *s* hash functions is defined for a sketch *S* of size *s*. Each sketch entry *S_i_* is defined as the minimum-valued *k*-mer after applying the hash function *Γ_i_* to all *k*-mers in the read. The resulting set of *s* minimum-valued *k*-mers, or minmers, comprise the sketch. Given a discrete *tf-idf* weight *w_q_* for each *k*-mer, we now modify the MinHash computation by applying *w_q_* hash functions |*Γ*_*i*,l_,…, *Γ_i,wq_*} per entry, rather than the single *Γ_i_* as before. For each sketch entry *S_i_*, the min-mer is then chosen as the minimum hash value computed across all functions. Because highly weighted *k*-mers are hashed more times, this increases the chance that they will be chosen as a min-mer. In order to properly match the same *k*-mers with different weights, we index *k*-mers using their fixed MurmurHash3 hash values, and the weighted values are only used to determine inclusion in the read sketches. The *tf-idf* approach replaces the previous approach, based on traditional all-or-nothing filtering of repetitive *k*-mers. We evaluated multiple scoring approaches including *tf-idf, idf* only (down-weighting common words), and no weighting on several bacterial and eukaryotic genomes. Both *tf-idf* and *idf* outperformed unweighted comparisons in terms of the resulting assembly continuity and accuracy, and were comparable to each other. We therefore utilize *tf-idf* by default due to its common use in the natural language field and other MinHash applications (Chum et al. 2008).

The updated MHAP version also implements bottom sketching for the second-stage filter (Ondov et al. 2016). In contrast to the first-stage filter, which uses multiple hash functions (Broder et al. 2000), bottom sketching uses a single hash function, from which the *s* minimum values are retained as the sketch (Broder 1997). The former approach has the advantage that the Jaccard similarity can be estimated for 1 versus *N* reads by a series of *s* hash table lookups. In bottom sketching, each comparison requires an *O(s)* merge operation, but as a benefit, any substring of the original string can be sketched by simply eliminating the min-mers from the original sketch that do not occur in the substring. For the bottom sketch, we now store a constant number of *k*-mers per read (default 1,500), and directly estimate the overlap error rate from these sketches using the Mash distance. The overlapping region is estimated as previously (Berlin et al. 2015), but also using the bottom sketch *k*-mers.

### Parallel Overlap Sort and Index

The downstream algorithms require efficient access to all overlaps for a single read, so the overlaps are organized using an indexed on-disk structure where all overlaps for a single read are listed sequentially. Canu parallelizes overlap computation into multiple jobs, each generating a compressed file of binary encoded overlaps and a file recording the number of overlaps for each read in that file. These files are combined into the master structure using a parallel bucket sort (Supplementary Figure S26). Since each read will have a different number of overlaps, and all overlaps for a given read must be in the same bucket in order for the bucket to be sorted, the number of overlaps per read is used to compute the ranges of reads assigned to each bucket. The size of a bucket is chosen such that each contains the same number of overlaps, and no bucket is larger than some specified maximum size. In parallel, each file of compressed overlaps is rewritten to a set of uniquely named buckets, and overlaps are duplicated and added to the appropriate bucket (e.g. read *A* overlaps *B*; and read *B* overlaps *A*). Note that as each input file creates its own set of buckets, no synchronization is needed between jobs. When all overlaps are copied into buckets, each bucket is loaded into memory, sorted, and output to a uniquely named file. Each bucket holds all of (and only) the overlaps for the range of assigned reads. Finally, an index describing the file and offset location for each read is created.

### Read Correction

Canu uses all-versus-all overlap information to correct individual reads. However, simply computing a consensus representation for each read using all overlaps could result in masking copy-specific repeat variants. Therefore, Canu uses two filtering steps to determine which overlaps should be selected to correct each individual read. The first is a global filter where each read chooses where it will supply correction evidence, and the second is a local filter where each read accepts or rejects the evidence supplied by other reads. This strategy attempts to overcome biases due to sequence quality and repeats. For example, reads with higher than average sequencing quality would tend to dominate the correction, regardless of if they were from the correct repeat copy. To prevent this, each read is only allowed to contribute to the correction of *C* other reads, where *C* is the expected read depth. The global filter scores each overlap (*overlap_length* * *identity*), and keeps only the *C* best overlaps for each read, thereby clustering repetitive reads with others likely to have originated from the same copy. When errors are uniformly distributed, we expect that reads are more likely to be grouped with reads from the same repeat copy, as they would have fewer total differences than reads from diverged repeat copies. A small fraction of mis-assigned reads is tolerable, as they will be outvoted during consensus correction. This strategy was first introduced by PBcR for the hierarchical correction and assembly of single-molecule reads (Koren et al. 2012). From this list, the local filter then selects the 2*C* best overlaps to each read for use in correction. The second filter is primarily a computational optimization.

Before computing the corrected sequence, the all-pair overlaps are used to predict the expected length of each read after correction (i.e. accounting for reads with partial or no overlaps). From these estimates, the longest reads up to a user-specified coverage depth are processed for correction. Corrected reads are generated using a modified implementation of the “falcon_sense” algorithm (Chin et al. 2016), which parallelizes the pairwise alignment step and removes and maximum read length limits. For a given read to be corrected, overlapping reads are aligned to it using Myers' O(ND) algorithm (Myers 1986). A directed acyclic graph (DAG) is created from the alignments, and the highest weight path is followed to generate a corrected sequence (Chin et al. 2016). Edges with weight less than four are omitted, which will split the original read when there is insufficient evidence for correction.

### Overlap Based Trimming

After correction, reads are trimmed by re-computing overlaps for the corrected reads and removing sequence that is not supported by other reads. The prior correction stage also trims low-coverage regions, but these initial overlaps are constructed without constructing a gapped alignment, which can result in imprecise trim points. When overlapping the corrected reads for trimming, a gapped alignment is computed for each overlap, and the trim points can be identified more precisely. Overlap-based trimming (OBT) was first described by Miller *et al*. (Miller et al. 2008) and Prüfer *et al*. (Prüfer et al. 2012), which focused on trimming Sanger, 454 and Illumina reads. Long reads with uniform error allow the algorithm to be simplified. Each read is trimmed to the largest portion covered to at least depth *C* by overlaps of at most *E* error and minimum length *L*. The parameters are technology specific and set to empirically derived defaults.

Once reads are trimmed, a second pass is made to detect any technology specific flaws, e.g. undetected hairpin adapters and chimeras (Eid et al. 2009; Jain et al. 2015). A hairpin adapter is detected by identifying when multiple reads have both forward and reverse overlaps around a common (short) sequence and there are few reads spanning this region. A chimeric junction is similarly detected by identifying a region with few, if any, spanning reads. In both cases, the original read is trimmed to the largest supported region.

### Overlap Error Adjustment

After trimming and before graph construction, Canu recomputes overlaps and makes a final attempt at detecting sequencing errors. This algorithm was first used in Holt *et al*. (Holt et al. 2002). The intuition is to improve separation between true sequencing differences (e.g. diverged repeats or haplotype) and false differences due to random sequencing error. Each read is corrected by a majority vote of its overlapping alignments, preserving differing bases only if there is sufficient support from other reads for this variation. The read sequence itself is not changed (doing so would invalidate the computed overlaps), but the reported error rate for each overlap is adjusted based on the alignment that would be generated had the sequencing errors been resolved. The algorithm requires two passes through the overlaps, the first pass detects probable sequencing errors in reads and the second pass applies those changes temporarily to reads to re-compute alignments and update the computed error rates.

### Graph Construction

The Bogart module builds an assembly graph using a variant of the “best overlap graph” strategy from (Miller et al. 2008). Overlaps are described as *containment*, if all bases in one read are aligned to another read, or *dovetail*, if involving only the ends of both reads. By definition, at least two read ends must be present in the alignment. A “best” overlap is the longest dovetail overlap to a given read end. Each read has two best overlaps, one on the 5′ end and one on the 3′ end. In the original method, best overlaps were picked from all overlaps up to a user supplied overlap error rate cutoff. In Bogart, best overlaps are picked after several filtering steps remove abnormally high-error overlaps, potential chimeric reads, and reads whose overlaps indicate a possible sequence anomaly. This results in a cleaner and more accurate graph construction.

After correction, trimming, and overlap error adjustment, all computed overlaps are used to pick an initial set of best edges. This set of best edges is used to compute the median and median absolute deviation (MAD) of the overlap error rate. This distribution represents the residual read error left after all prior corrections, and a low average overlap error rate cutoff indicates good sequencing data and successful correction. A maximum overlap error rate cutoff is automatically computed from this distribution as six MADs away from the median, and overlaps with an error greater than this cutoff are not used during graph construction. This cutoff, which is typically less than 2% for good PacBio data (average median 0.232% and average MAD 0.138% for PacBio datasets in this paper), determines the ability of the algorithm to separate closely related repeats and haplotypes.

In addition to filtering poor overlaps, Bogart filters suspicious reads that may have evaded proper trimming and correction. First, reads that are not fully covered by overlaps below the overlap error rate cutoff are flagged as potentially chimeric and excluded from graph construction. Second, best overlaps are usually mutual, i.e. the best overlap from *A* is to *B* and the best overlap from *B* is to *A*. For a pair of reads, non-mutual best overlaps are often caused by Indels, making the overlap length slightly longer or shorter compared to the mutual best overlap. Thus, reads with a large overlap size difference are also excluded (Supplementary Figure S27).

The resulting set of reads and best overlaps define the best overlap graph. Initial contigs are then constructed from the best overlap graph as in (Miller et al. 2008), and an error rate profile is generated for each contig from the error rate of overlaps used to build it. A median and MAD value is computed for each window in the contig based on the overlaps falling in it to generate an error profile. This error profile is recomputed after each phase of the algorithm, and is used to determine if external reads have valid overlaps to the contig.

Bogart next attempts to include contained (Fasulo et al. 2002) and previously filtered reads into the contigs. All overlaps to these reads are used to compute a set of potential contig placements, scored by the average overlap error rate. If this average error rate exceeds the pre-computed error profile for the contig region the read is likely from a diverged repeat or a heterozygous variant, and the placement is rejected. The placement with the lowest average error is accepted, and the read is placed. This strategy differs from the original strategy from (Miller et al. 2008) that placed contained reads based on the highest quality containment overlap, which could incorrectly place a read when the true location had no containing read. Reads that remain unplaced after this phase are output as “unassembled.”

An assembly bubble occurs when there is more than one reconstruction of a specific locus caused by haplotype differences (Fasulo et al. 2002; Zerbino and Birney 2008; Koren et al. 2011; Nijkamp et al. 2013; Chin et al. 2016). Small differences, tens of base pairs in size, are typically not detectable from overlaps alone because the difference is insignificant compared to the size of the overlap. Larger differences can result in two, mostly redundant, contigs covering the same locus. The haplotype with more reads is often reconstructed in a large contig spanning the locus, and the haplotype with fewer reads as just the variant region (the bubble). Currently, contigs with fewer than a minimum threshold of reads, or with more than 75% of the reads with an overlap to some other contig, are considered potential bubbles. Reads in these contigs are then placed, using the mechanism for placing unplaced reads as above, into all other contigs where possible using heuristics. Improved mechanisms for resolving bubbles within the assembly graph, and ultimately producing a fully phased assembly, is an area of ongoing research and left for future work.

Despite careful filtering, the greedy construction algorithm remains prone to error and the graph will be missing edges compared to a full string graph representation, so a final step is required to add missing edges and break incorrectly assembled contigs. Using the all-pairs overlap information, every assembled contig is annotated with compatible read placements, again using the read placement mechanism and all reads from non-bubble contigs. Only overlaps that meet the global and local contig error rate thresholds are considered. The resulting annotated regions indicate alternative branch points in the full overlap graph, and a correct contig reconstruction is confirmed by the presence of spanning reads or overlaps. Unresolved regions are marked as repeats, the contig is split, and additional edges are added to form the final assembly graph.

### Contig Consensus

Canu generates a consensus sequence for each contig using a modified version of the “pbdagcon” algorithm (Chin et al. 2013). Briefly, a template sequence is constructed for each contig by splicing reads together from approximate positions based on the best overlap path. This template is accurate within individual reads, as they have previously been error-corrected, but may have Indel errors at read boundaries due to inaccuracy in the overlap positions. To correct this, all reads in the contig are aligned to the template sequence in parallel using Myers' O(ND) algorithm (Myers 1986) and added to a DAG. The DAG is then used to call a consensus sequence as in (Chin et al. 2013).

### Assembler Versions

Falcon v0.4.1 as of 2016-03-16 (commit c602aad3667b3fd49263028dac44da8e42caa 17c). Minimap/miniasm as of 2016-03-16 (commit 1cd6ae3bc7c7a6f9e7c03c0b7a93a12647bba244 minimap, 17d5bd12290e0e8a48a5df5afaeaef4d171aa133 miniasm). SPAdes v3.7.1. Canu v1.3 (Supplementary Note 5).

### Data Access

The *Bacillus anthracis* Sterne 34F2 sequencing data has been deposited at NCBI under BioProject PRJNA357857 and the *Yersinia pestis* 195/P sequencing data under PRJNA357858. All other sequencing was obtained from external sources and listed in Supplementary Note 2.

## Acknowledgements

We thank Celera and Pacific Biosciences for open source software that was critical for the development of Canu, and also John Urban and all other Canu users who provided early testing and feedback on the software. We thank Shaun Jackman and the other reviewers for their considered reviews, and one anonymous reviewer for providing a motivating example on repeat separation. This research was supported in part by the Intramural Research Program of the National Human Genome Research Institute, National Institutes of Health, and under Contract No. HSHQDC-07-C-00020 awarded by the Department of Homeland Security (DHS) Science and Technology Directorate (S&T) for the management and operation of the National Biodefense Analysis and Countermeasures Center (NBACC), a Federally Funded Research and Development Center. The views and conclusions contained in this document are those of the authors and should not be interpreted as necessarily representing the official policies, either expressed or implied, of the DHS or S&T. In no event shall the DHS, NBACC, S&T or Battelle National Biodefense Institute (BNBI) have any responsibility or liability for any use, misuse, inability to use, or reliance upon the information contained herein. DHS does not endorse any products or commercial services mentioned in this publication. This work utilized the computational resources of the NIH HPC Biowulf cluster (http://hpc.nih.gov).

